# Rod-cone signal interference in the retina shapes perception in primates

**DOI:** 10.1101/825851

**Authors:** Adree Songco-Aguas, William N Grimes, Fred Rieke

**Author notes:** Equal contributions (Adree Songco-Aguas, William N Grimes). Lead contact: Fred Rieke.

## Abstract

Linking the activity of neurons, circuits and synapses to human behavior is a fundamental goal of neuroscience. Meeting this goal is challenging, in part because behavior, particularly perception, often masks the complexity of the underlying neural circuits, and in part because of the significant behavioral differences between primates and animals like mice and flies in which genetic manipulations are relatively common. Here we relate circuit-level processing of rod and cone signals in the non-human primate retina to a known break in the normal seamlessness of human vision – a surprising inability to see high contrast flickering lights under specific conditions. We use electrophysiological recordings and perceptual experiments to identify key mechanisms that shape the retinal integration of rod- and cone-generated retinal signals. We incorporate these mechanistic insights into a predictive model that accurately captures the cancellation of rod- and cone-mediated responses and can explain the perceptual insensitivity to flicker.

## Introduction

Computation in neural circuits often relies on the distinct processing of input signals within different parallel pathways and the control of circuit outputs by the convergence and integration of these parallel signals. Although these computational processes recur throughout the central nervous system, there are very few circuits in which we know enough about which cell types belong to which pathways to relate mechanisms to circuit function, much less to behavior. Here we study retinal signaling and perception under conditions in which behavior depends on a combination of signals originating in the rod and cone photoreceptors. Under these conditions, interactions between rod and cone signals shape multiple aspects of visual perception (reviewed by (Buck, 2004; Buck, 2014; Grimes, Songco-Aguas, & Rieke, 2018b)). Investigating the origin of these interactions provides a rare opportunity to relate the mechanisms governing parallel processing directly to perception.

Several common motifs shape parallel processing in neural circuits (reviewed by (Cajal, 1937; Dunn & Wong, 2014; Braganza & Beck, 2018)). First, input signals can diverge to distinct parallel pathways, which then process those common inputs differently. Common inputs, for example, typically diverge to excitatory and inhibitory subcircuits with distinct properties such as kinetics. Second, outputs of several parallel pathways can converge onto a target neuron or population of target neurons to control circuit outputs. Integration of excitatory and inhibitory synaptic inputs is a ubiquitous example of this motif. Third, interactions between parallel pathways can shape the signals that they convey. Lateral inhibition provides a common example of this final motif. Combinations of these motifs underlie many computational properties of neural circuits (reviewed by (Isaacson & Scanziani, 2011; Jadzinsky & Baccus, 2013)), including sharpening neural tuning, governing selectivity of different parallel circuit outputs for specific stimulus features, and separating signals of interest from other ‘background’ signals or noise.

The specific circuit properties responsible for sensory behavior are often obscured by the seamless nature with which we perceive the world. Breaks in this seamlessness can provide a window into the underlying mechanisms. Such an approach, for example, links nonlinear distortions produced in the inner ear to auditory perception (Barral & Martin, 2012; Jaramillo, Markin, & Hudspeth, 1993). Optical illusions are another example (Eagleman, 2001), although many are sufficiently complex that they likely rely on mechanisms located in multiple visual areas. Here, we focus on a surprising break in the seamlessness of visual perception: the perceptual insensitivity to high contrast flickering lights under conditions where both rods and cones are active (MacLeod, 1972). The requirement for coactivation of rods and cones and the dependence of this effect on the specific temporal frequency of the stimulus suggest a destructive interference between rod- and cone-mediated signals. Because rod and cone signals converge within the retina to modulate the responses of common retinal output cells, such destructive interference is likely to occur within retinal circuits. Indeed, destructive interference of rod and cone signals is apparent in electroretinograms (McAnany, Park, & Cao, 2015), although such measurements do not identify the mechanism or location of such interference.

Here, we test whether rod and cone signals interfere in the responses of retinal output neurons and the implications of such interference for the mechanistic operation of retinal circuits under conditions in which both rods and cones contribute to vision (i.e. mesopic conditions such as moonlight). We first reproduce the perceptual interference between flickering rod and cone signals seen previously (MacLeod, 1972). We then use similar stimuli to probe retinal output signals and find that rod and cone signals indeed destructively interfere within the retina. These experiments directly reveal the kinetic differences between rod and cone signals underlying this destructive interference. Finally, we use our physiological results to construct a computational model that can account for the destructive flicker interactions and predict other temporal interactions between rod and cone signals. Together, these results link a clear and unexpected perceptual result to parallel processing within retinal circuits.

## Results

To connect human perception and retinal processing, we performed parallel non-human primate electrophysiological experiments and human psychophysical experiments. Using the same stimulus conditions, we identified 1) retinal circuits that exhibit similar interactions to those observed perceptually, 2) signal properties that were necessary for the interactions, and 3) manipulations that make predictive changes to the interactions that could be tested experimentally.

### Rod-cone signal interference in human perception

We used several psychophysical tasks to probe perceptual interactions between time-varying rod and cone signals. These tasks relied on the ability to preferentially activate rod photoreceptors with dim short-wavelength light and long-wavelength sensitive (L) cones with long-wavelength light (see Methods and (Grimes et al., 2015).

Tasks #1 and #2 measured independent thresholds for rod- and cone-preferring flicker. After dark-adapting for 20 minutes, a rod- or cone-preferring stimulus was modulated sinusoidally in time in an observer’s peripheral visual field (2° spot at 10° eccentricity, 2 s duration). After each stimulus presentation, observers indicated whether they could detect the flickering of the spot (Figure 1A). The results were recorded and used to update the spot contrast (i.e. the change in luminance as a percentage of the mean) for the next presentation. This process was repeated multiple times to estimate perceptual flicker detection thresholds (see Methods). Blocks of rod- and cone-preferring stimuli were interleaved and analyzed separately. We repeated this process for a range of stimulus frequencies (4-9.5 Hz) to extract perceptual thresholds for isolated rod and cone flicker (Figure 1B). The threshold for rod flicker depended more strongly on frequency than that for cone flicker, as expected from previous work and the more rapid kinetics of cone-mediated responses (Lee, Smith, Pokorny, & Kremers, 1997; Verweij, Dacey, Peterson, & Buck, 1999).

**Figure 1.**
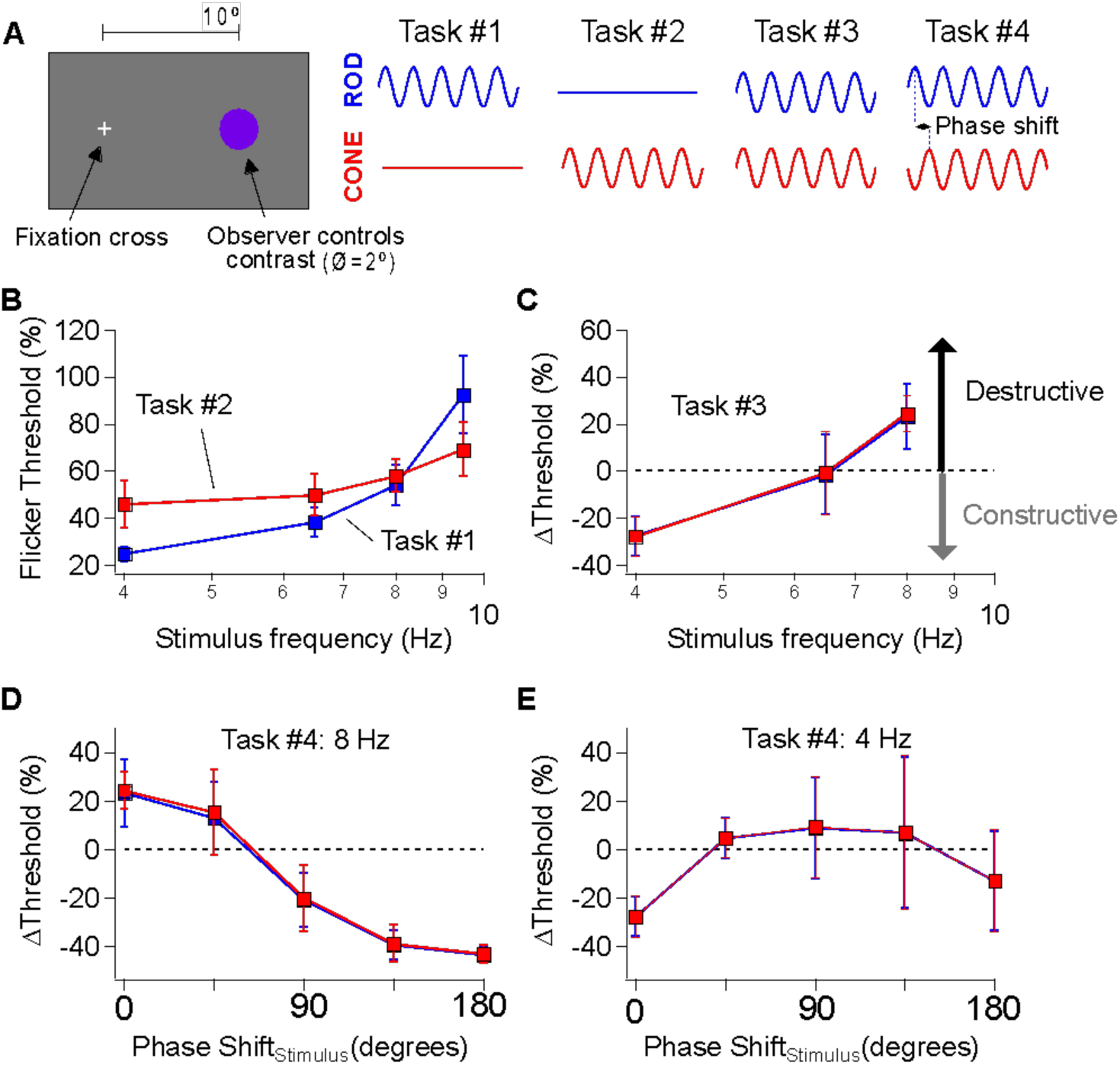
Interference between rod- and cone-generated signals in human perception. A) General setup for human perceptual experiments to probe rod-cone signal integration. B-C) Mean flicker thresholds for rod- (B, blue) and cone- (C, red) isolating stimuli across a range of temporal frequencies. Purple data points reflect thresholds obtained from trial when both rod and cone stimuli flickered simultaneously. D-E) Introducing phase shifts between rod and cone flicker (stimuli) shift interactions from destructive to constructive (e.g. 8 Hz; D) and vice-versa (e.g. 4 Hz; E).

Task #3 measured combined thresholds for rod- and cone-preferring flicker. We presented observers with simultaneously flickered rod- and cone-preferring stimuli at a fixed contrast ratio set by the thresholds for detection identified in Tasks #1 and #2 (see Methods). Observers repeated the threshold measurements described for Tasks #1 and #2 using these combined stimuli. We then determined the change in threshold for the combined stimuli compared to that of the independent rod and cone stimuli; a change in threshold of 0% means that the combined flickering stimuli were just detectable when the constituent rod and cone stimuli were at their individual thresholds. Perceptual thresholds for combined stimuli at frequencies ≤6.5 Hz were reduced compared to trials in which rod- and cone-preferring stimuli were flickered separately, whereas thresholds for combined stimuli at 8 Hz were increased (Figure 1C). These results are consistent with the perceptual experiments originally conducted by Don MacLeod (MacLeod, 1972), who hypothesized that shifts in flicker thresholds reflect constructive and destructive interference between rod and cone signals. Following this logic, the mode of signal interference (i.e. constructive or destructive) depends on the phase shift between responses to the rod- and cone-preferring stimuli; this phase shift in turn is determined by the delay between rod and cone signals produced within the associated neural circuits and the frequency of the stimulus.

Task #4 determined how thresholds for combined stimuli were affected by an added phase shift between the rod- and cone-preferring stimuli. The destructive signal interference hypothesis is based on the slower kinetics of rod signals compared to cone signals. If this hypothesis is correct, shifting the timing (or equivalently phase) of rod-preferring stimuli relative to cone-preferring stimuli should modify rod-cone interactions (e.g. shift them from destructive to constructive) as the relative timing of rod and cone signals changes. Indeed, at 8 Hz, introducing a phase shift between the rod and cone stimuli substantially lowered perceptual thresholds compared to trials with zero phase shift (Figure 1D). Conversely, introducing a phase shift between 4 Hz rod and cone flicker substantially increased flicker thresholds (Figure 1E), although the dependence of threshold on the added phase shift was smaller than that observed for 8 Hz flicker. The systematic dependence of threshold on the relative phases of the rod and cone flicker provides additional evidence that perceptual thresholds reflect constructive and destructive interference between kinetically-distinct rod and cone signals.

### Rod-cone signal interference in the retinal outputs

Rod- and cone-mediated signals converge within retinal circuits to modulate spike responses of a common set of retinal ganglion cells (Figure 2A; (Enroth-Cugell, Hertz, & Lennie, 1977; Gouras & Link, 1966)). Hence any visual area receiving input from the retina, including any area involved in the perceptual phenomena illustrated in Figure 1, receives intermixed rod- and cone-mediated signals. This anatomical and functional convergence predicts that rod-cone flicker interference might occur within the retina (MacLeod, 1972), and recent *in vivo* ERG experiments support this hypothesis (McAnany et al., 2015).

**Figure 2.**
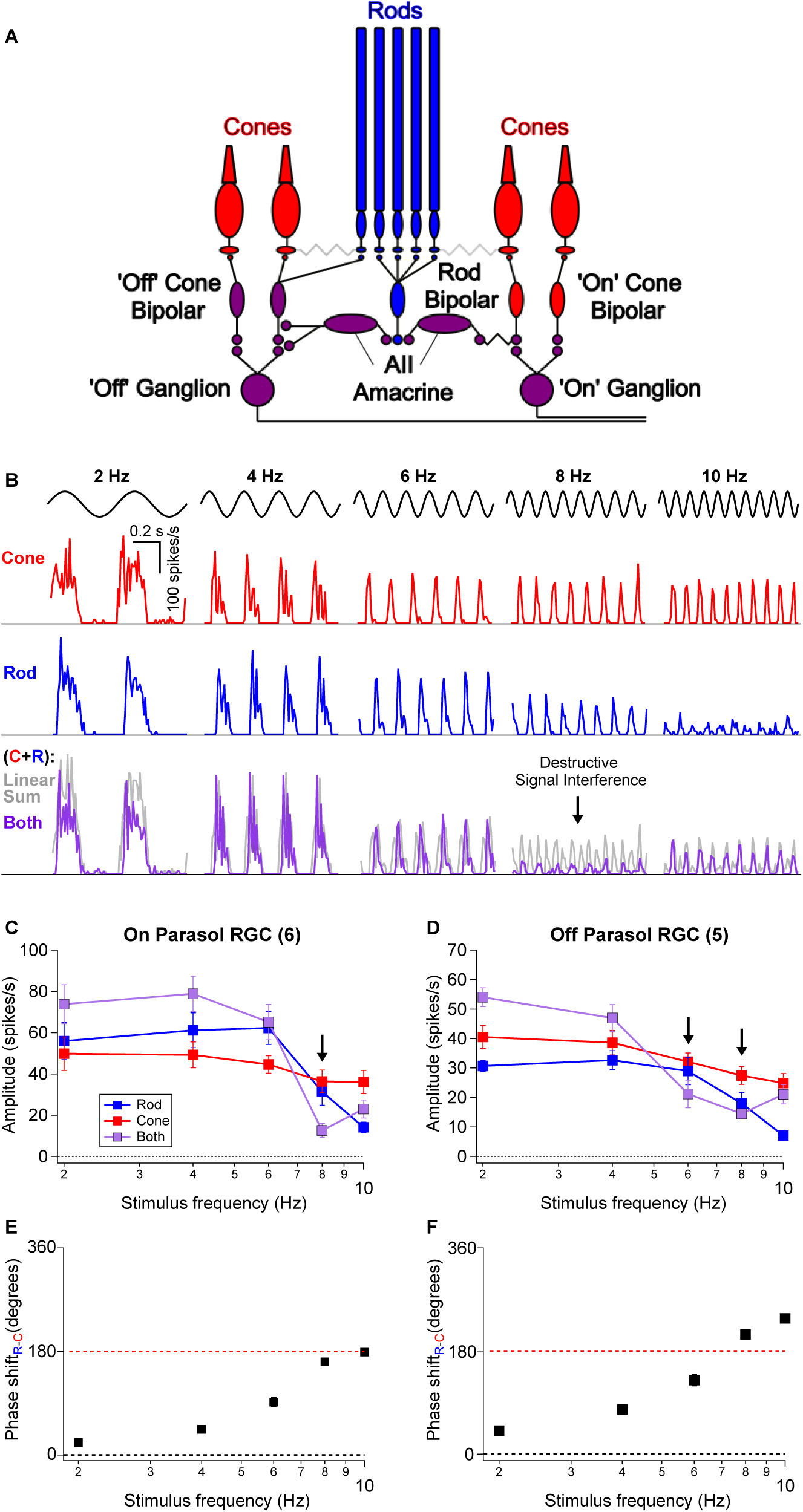
Interference between rod- and cone-generated signals in isolated non-human primate retina. A) Diagram of the retinal circuits that convey rod and cone signals to ‘On’ and ‘Off’ retinal ganglion cells. Rod signals can be transmitted through multiple routes, but recent work (Grimes et al., 2018 eLife) indicates that the dedicated rod bipolar pathway is the primary conduit in primates. Rod bipolar cells initially excite AII amacrine cells This in turn excites ‘On’ ganglion cells through gap junctions with presynaptic ‘On’ cone bipolar cells and inhibits ‘Off’ ganglion cells through both direct inhibition to the ganglion cell’s dendrites and inhibition of presynaptic ‘Off’ cone bipolar cells. Because signals from L- and M-cones are transmitted to ganglion cells through ‘On’ and ‘Off’ cone bipolar cells, rod-cone signal integration largely occurs within the axons of these bipolar cells. B) Spike responses to cone (*top*, red), rod (*middle*, blue), or combined rod-cone (*bottom*, purple) flicker (2-10 Hz) in an On Parasol retinal ganglion cell. A strong suppressive interaction was observed when rod and cone stimuli were flickered together at 8 Hz. C-D) Mean spike responses (sinusoidal fit, see Methods) as a function of stimulus frequency for 6 On Parasol RGC recordings (C) and 5 Off Parasol RGC recordings (D). E-F) Relative timing differences (i.e phase shifts) between isolated rod and cone responses for On (E) and Off (F) Parasol RGCs. Markers and error bars in C-F represent mean±SEM.

To directly test this proposal, we recorded the spiking activity of parasol RGCs from dark-adapted isolated primate retinas (see Methods) in response to rod- and cone-preferring stimuli like those used in Figure 1. For each experiment, we began by independently adjusting the contrasts of rod and cone stimuli (flickered at 8 Hz) to produce similar levels of spiking activity in On Parasol RGCs. These contrasts were then held constant as we probed responses of On and Off parasol RGCs across a range of temporal frequencies (2-10 Hz). On Parasol RGCs showed robust periodic spiking activity in response to cone flicker across all temporal frequencies tested (Figure 2B *top row*). Rod flicker also produced robust periodic activity in the same cells for frequencies below 10 Hz (Figure 2B *middle row*).

On Parasol RGCs responded strongly to joint rod/cone flicker at stimulus frequencies ≤6 Hz and ≥10 Hz, but showed little modulation at a stimulus frequency of 8 Hz (Figure 2B *bottom row*, Figure 2C). The gray lines in Figure 2B show the linear sum of the responses to rod and cone stimuli delivered individually. At low frequencies, responses to the joint stimuli fell short of this linear sum, likely due to saturation of the firing rate (the peak firing rate to the joint stimuli often exceeded 300 spikes/s). Responses to the joint stimuli were also much smaller than the linear sum of the individual responses at 8 Hz; Off parasol RGCs showed a similar frequency-dependent response suppression at both 6 and 8 Hz flicker (Figure 2D). Saturation of the firing rate cannot explain this nonlinear interaction given the low firing rates. Instead, the small response at 8 Hz suggests a destructive signal interference like that needed to account for the perceptual results of Figure 1.

A closer inspection of the time course of responses to independent flicker revealed a lag in rod signals relative to cone signals; this lag became larger relative to the period of the stimulus for higher frequencies. We divided the relative delays by the stimulus period to estimate the phase shift between individual rod and cone responses (Figure 2E-F). This analysis indicates that at 8 Hz rod and cone signals are near-perfectly (i.e. 180 degrees) out of phase. Hence the relative timing of rod and cone signals in the retinal output is in close agreement with that inferred from perceptual flicker cancellation (Figure 1 and (MacLeod, 1972)).

If rod-cone signal interference depends on the relative timing of retinal signals, then we should be able to shift interactions from destructive to constructive and vice versa by introducing a relative delay between the rod and cone stimuli (as in the perceptual task of Figure 1D-E). At 8 Hz with zero phase shift (i.e. no delay), simultaneous presentation of rod and cone flicker produced only weak activity in RGCs (Figure 3A *left*). The introduction of a phase shift between the 8 Hz flickering stimuli was sufficient to shift rod-cone interactions from destructive to constructive (Figure 3A right, Figure 3B). Conversely, at 4 Hz with zero phase shift, simultaneous presentation of rod and cone flicker produced activity in RGCs that was greater in amplitude than responses to rod or cone flicker alone (Figure 3C *left*). Introduction of a phase shift to the 4 Hz stimuli shifted the rod-cone interactions from constructive to destructive (Figure 3C right, Figure 3D).

**Figure 3.**
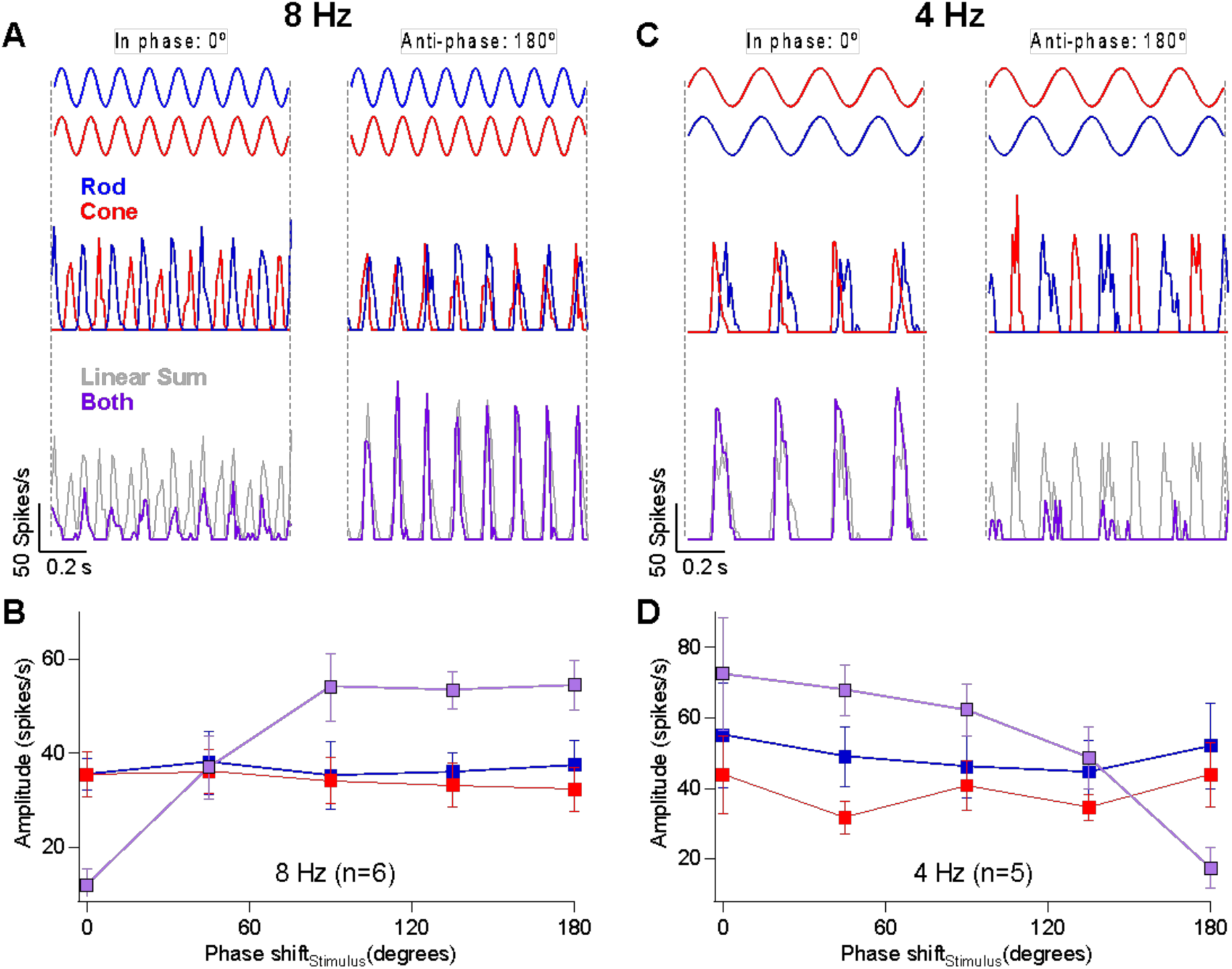
Rod-cone interference can be constructive or destructive depending on the relative timing of the retinal responses. A) Responses of an On Parasol RGC to 8 Hz flicker when the rod and cone stimuli are in-phase (*left*) or anti-phase (*right*). B) Response amplitudes collected from 6 On Parasol RGCs to rod, cone or combined flicker for a range of phase shifts between the stimuli. Responses to 8 Hz flicker shift from constructive to destructive after introduction of a phase shift. C-D) The same demonstration as **A-B**, but this time using 4 Hz flicker. Responses to 4 Hz flicker shift from destructive to constructive after introduction of a phase shift.

Together, these results demonstrate that constructive and destructive interference of rod and cone signals within retinal circuits substantially shape signals sent to the magnocellular layers of the LGN. Furthermore, the dependence of these signal interactions on stimulus frequency and the relative timing of rod- and cone-preferring stimuli closely match those observed perceptually.

### Absolute signal delays depend on stimulus frequency

Previous attempts to model perceptual rod-cone flicker interactions assume summation of rod and cone signals with a *fixed* absolute delay between the signals; this delay is often assumed to be half the period of the stimulus frequency which maximizes cancellation (e.g., cancellation at 8 Hz corresponds to a 62.5 ms delay). Furthermore, this delay is assumed to arise from the slower light responses of rods compared to cones and the additional synapses involved in conveying rod signals through the retina (Figure 2A; (Sharpe, Stockman, & MacLeod, 1989)). Our direct recordings from On parasol RGCs, however, indicate that the delay of rod signals relative to cone signals is not fixed but instead depends on stimulus frequency (Figure 4).

**Figure 4.**
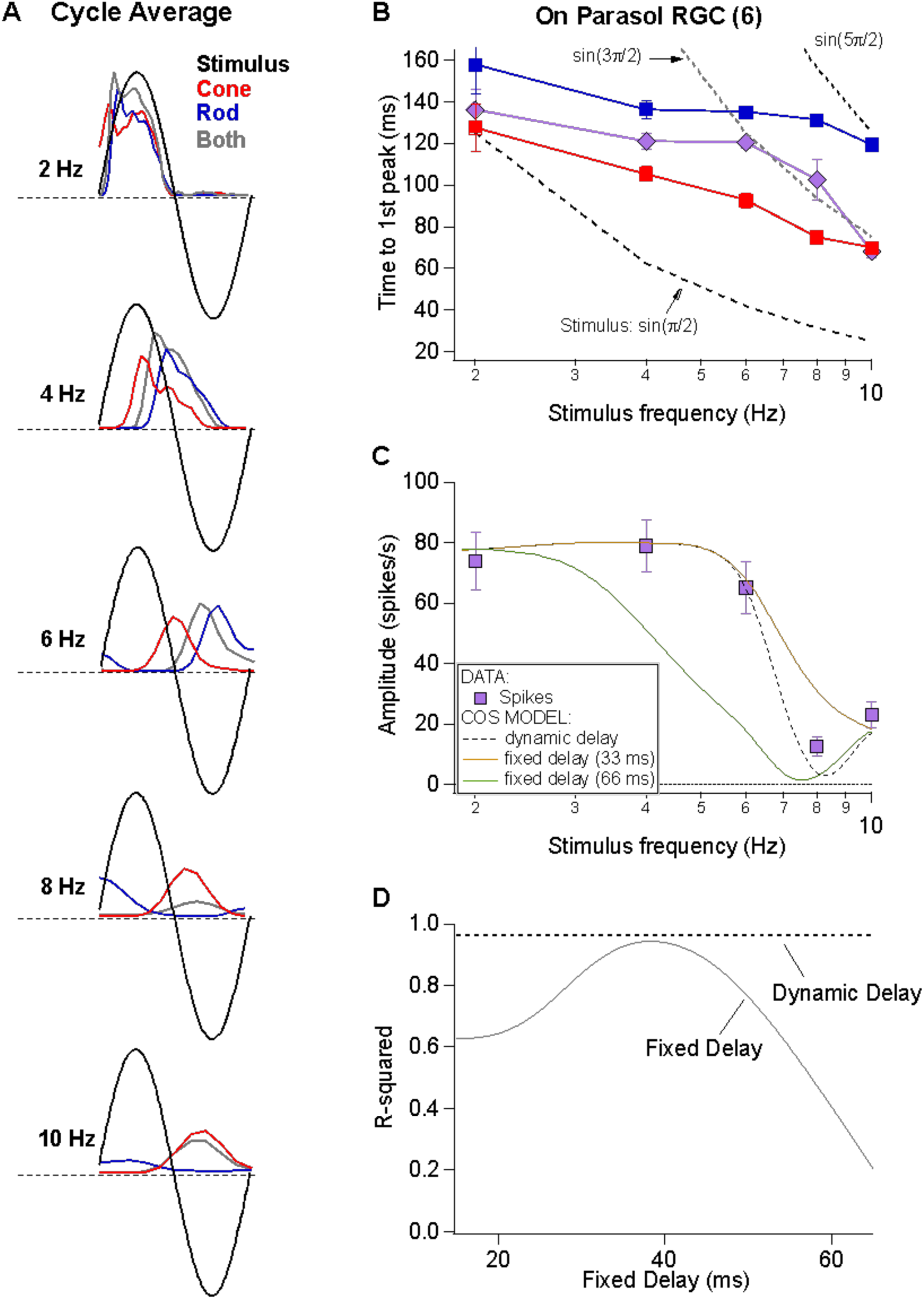
Absolute timing data and vector summation model. A) Time to first peak of the response to rod, cone or simultaneous modulation of spot contrast across frequency. Dashed lines represent the timing of the peaks and troughs of the sine wave stimuli. B) Fitting measured data with a vector summation model. Empirically measured dynamic delays provide better estimates than fixed delays. D) Goodness of fit for models using a range of fixed delays and an empirically-measured dynamic delay.

Cycle-averaged responses to rod, cone and combined flicker illustrate the relative timing of the retinal responses (Figure 4A). With increasing frequency, the delay of both rod- and cone-mediated responses relative to the stimulus increased and the separation between rod- and cone-mediated responses increased. Figure 4B summarizes these empirical measurements of response timing across cells and temporal frequency. The fixed-delay picture predicts that the vertical separation of the rod and cone traces remains constant. Instead, the data show larger rod vs cone signal delays at higher temporal frequencies (e.g. 2 Hz: 30±9 ms vs. 8 Hz: 56±5 ms, p=0.003, n = 6).

To test the impact of these frequency-dependent delays on the integration of rod and cone signals, we used a simple summation model in which we added sinusoidal responses, phase shifted to reflect either measured or assumed relative delays of rod and cone responses. The solid lines in Figure 4C show the results of this modeling for *fixed* rod-cone delays of 33 and 66 ms, approximating the relative delays often assumed for the fast and slow rod pathways (Kilavik & Kremers, 2006; Sharpe et al., 1989); the dashed line shows results for a dynamic delay model in which the delays have a frequency dependence taken directly from Figure 4B. Models incorporating a fixed delay were unable to account for the responses to the full range of frequencies tested. Models with a 33 ms fixed delay correctly predicted combined responses to low (e.g. 2-6 Hz) and high (e.g. 10 Hz) frequency stimuli (R^2^=0.9 for the full range of frequencies tested), but failed to accurately predict the observed cancellation frequency (Figure 4C). Models with a 66 ms fixed delay more accurately captured the cancellation frequency, but performed poorly at low frequencies (Figure 4C,D, R^2^=0.17). Models incorporating the measured frequency-dependent delays provided good predictions for the full range of frequencies tested (Figure 4C,D,R^2^=0.96 across frequencies).

These results argue that delays between rod and cone signals reaching ON parasol RGCs are not fixed, but instead depend dynamically on stimulus frequency. This is an important departure from the fixed-delay models often used to interpret psychophysical findings (see Discussion).

### Rod-cone signal interference in the RGC excitatory synaptic inputs

Destructive rod-cone interactions in the retinal output could depend on a number of neural mechanisms, including the integration of excitatory and inhibitory synaptic inputs in the ganglion cells. To identify the origins of rod-cone signal interference, we isolated and recorded excitatory and inhibitory synaptic input to On Parasol RGCs using the whole-cell voltage-clamp technique (see Methods). These recordings revealed destructive interference within the excitatory (Figure 5A-B) and inhibitory (Figure 5C) inputs to On Parasol RGCs at 8 Hz. Excitatory (Figure 5D) and inhibitory (Figure 5E) synaptic inputs exhibited frequency-dependent phase shifts similar to those observed in spike recordings (Figure 2E). The similarity of the phase shifts for excitatory inputs and spike responses indicates that the dynamic delays explored in Figure 4 are a property of retinal circuits rather than an intrinsic property of the ganglion cells.

**Figure 5.**
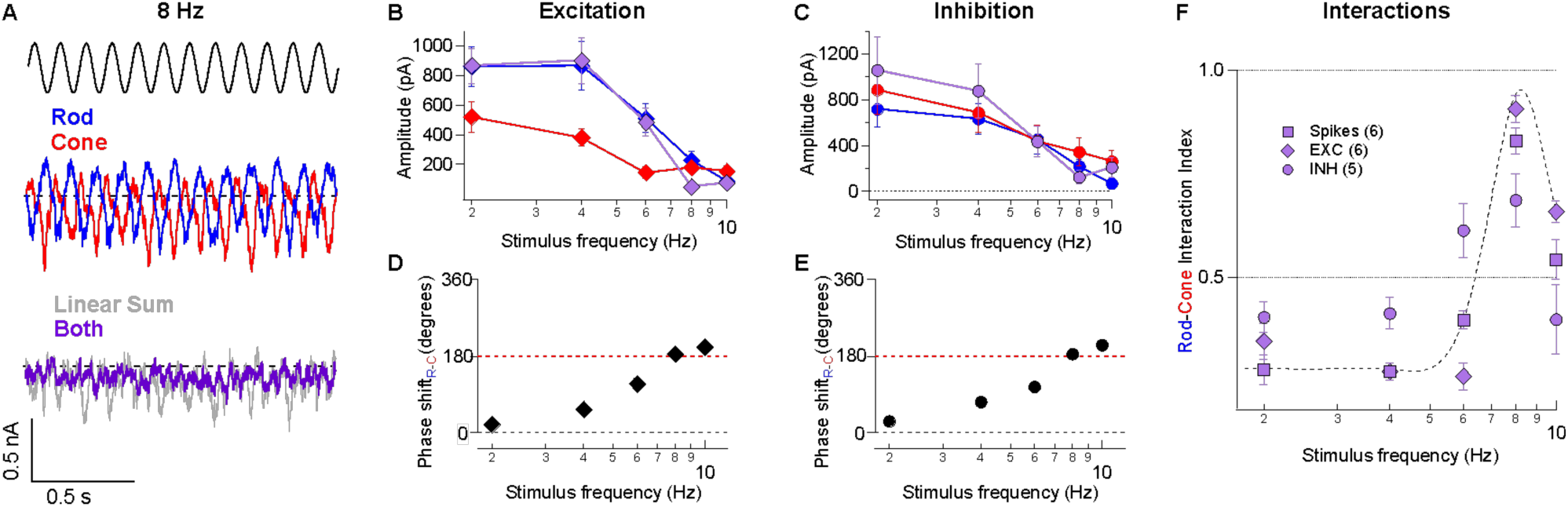
Rod-cone interference is present in the excitatory and inhibitory synaptic inputs to RGCs. A) Voltage-clamp recordings of excitatory synaptic input to an On Parasol RGC in response to 8 Hz rod (blue trace), cone (red trace), or combined (rod-cone) flicker (grey trace). B-C) Excitatory (B) and inhibitory (C) synaptic response amplitudes (sinusoidal fit, see Methods) for stimulus frequencies ranging 2-10 Hz for 6 On Parasol RGC recordings. D) Relative timing differences (i.e phase shifts) between isolated rod and cone excitatory inputs to On Parasol RGCs. E) Same as in D but for inhibitory synaptic input. F) Plot of the rod-cone nonlinear interaction index (see Methods) as a function of stimulus frequency. Recordings of excitatory and inhibitory synaptic input reveal a similar degree of destructive interference to that of spike recordings. Markers and error bars in B-F represents mean±SEM.

As is the case for the spike outputs, the linear sum of the responses to the separate rod and cone stimuli failed to predict responses to the joint stimuli (Figure 5A). This failure is consistent with integration of rod and cone signals in On pathways prior to a rectifying nonlinearity, likely at the output synapse of the presynaptic On cone bipolar cells (Figure 2A). We quantified the rod-cone interaction using a nonlinear interaction index (*II*, see Methods), where a value of 1 represents complete suppression and a value of zero reflects a simple linear sum. The *II* for spikes, excitatory synaptic input and inhibitory synaptic input at 8 Hz was 0.83 +/− 0.03 (n=6), 0.91 +/− 0.03 (n=6), and 0.69 +/− 0.06 (n=5) (Figure 5F). These results argue that the destructive interference observed at 8 Hz in the RGC spike output is largely inherited from the integration of rod and cone signals in upstream retinal circuits.

### Predictive model for rod-cone signal integration in the retina

Sensory networks encounter a steady stream of temporally- and spatially-varying information, and a true understanding of a given system includes the ability to predict the system’s response across a broad range of stimuli. With this long-term goal in mind, we developed a kinetic model that can predict retinal responses to several time-varying rod and cone stimuli, including interactions between them (Grimes et al., 2015).

Our model of retinal integration is rooted in the framework of linear-nonlinear cascade models (Chichilnisky, 2001). This general class of computational models is composed of two empirically-derived elements (Figure 6B): 1) a linear filter that accounts for the response kinetics, and 2) a static or time-independent nonlinearity that accounts for properties such as rectification at synapses or in spike generation. To estimate these model elements, rod- or cone-preferring spatially-uniform gaussian noise (cutoff frequency = 40 Hz; Figure 6A) was delivered to the retina while recording excitatory synaptic input to On Parasol RGCs. Both stimuli elicited RGC responses of comparable strength. The relation between the stimulus and response was used to extract linear filters and static nonlinearities for both rod and cone stimuli (Figure 6B, right; see Methods). As previously observed (Grimes et al., 2015), filters derived using rod stimuli were slower and typically more biphasic than filters derived for cone stimuli, whereas rod- and cone-derived nonlinearities had similar shapes. To verify the accuracy of our LN models, we tested whether predictions of single rod or cone models matched the measured excitatory synaptic inputs in response to the corresponding rod or cone stimuli (Figure 6C). Across cells, rod and cone models performed similarly well, with an average fraction of variance explained of ∼0.8 (Figure 6D).

**Figure 6.**
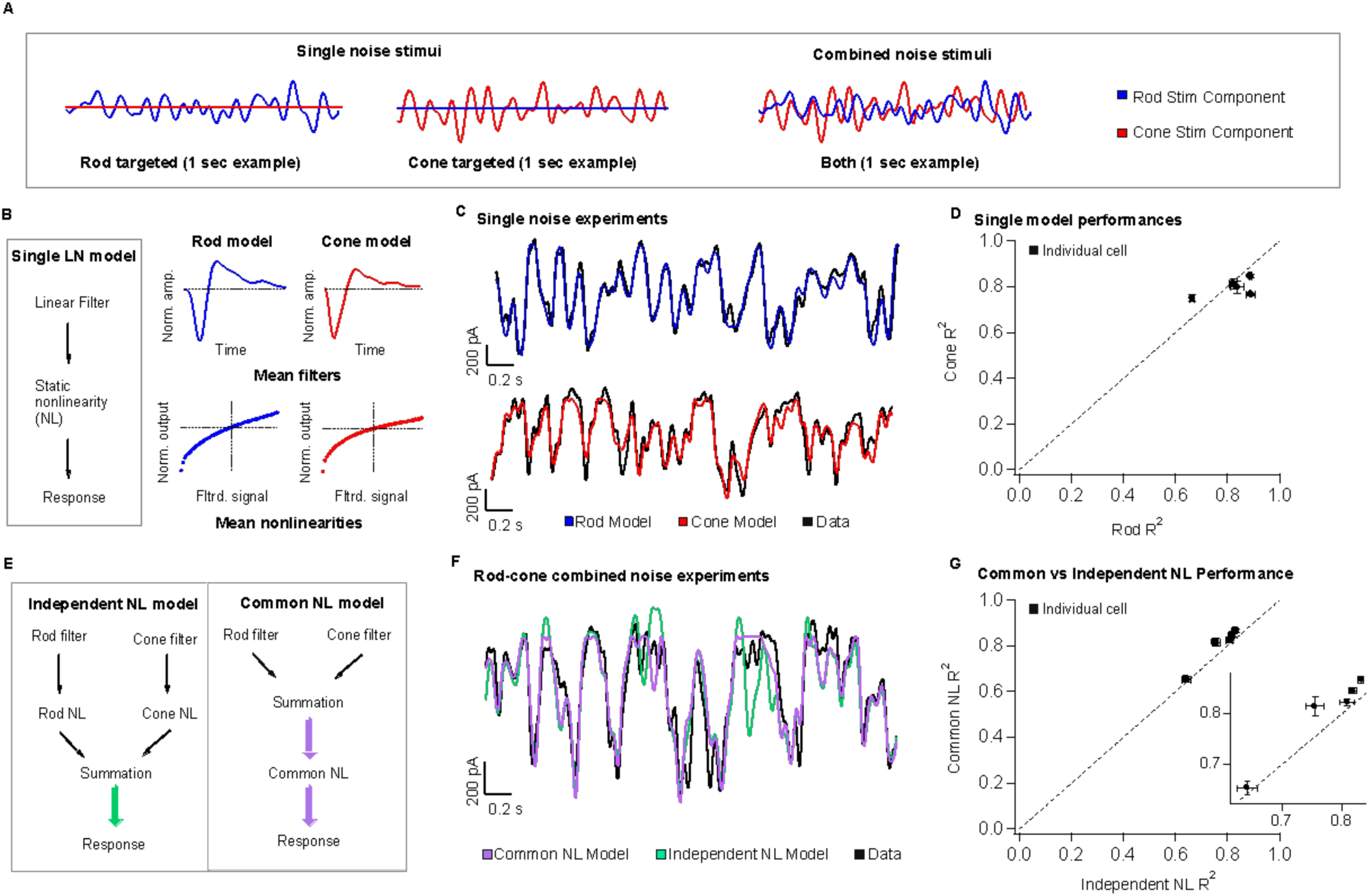
A simple linear-nonlinear model supports pre-synaptic integration of rod and cone signals. A) *left* The model architecture for a single-pathway model – a linear filter stage followed by a nonlinear transformation. *right* Each stage of the model is trained on excitatory current data from On Parasol RGCs, in response to rod and cone targeted white noise stimuli. Presented here are the mean single model fits and variance across 5 different cells. B) Example data traces, with the model predictions overlaid. C) The single model performances across five cells is presented as the measure of explained variance of the model. D) Two different three-stage model architectures are presented. On the left, the two branches of the rod-cone model integrate after independent nonlinear transformations, corresponding with exclusively post-synaptic integration. On the right, the two branches integrate before a common nonlinear transformation, corresponding with exclusively pre-synaptic integration. E) *top* Uncorrelated rod and cone targeted noise stimuli are presented simultaneously in experiments to find the best model architecture. *Bottom* Example trace, qualitatively demonstrating the common nonlinearity model outperforming the independent nonlinearity model. (Supplementary Figure 3 shows a performance breakdown across different rod and cone contributions to the final signal.) F) Quantification of common nonlinearity model outperformance across all five cells, as measured by the explained variance.

The individual components of the rod and cone LN models were used to construct a single model that predicts excitatory synaptic currents in response to simultaneous time-varying rod and cone stimuli (Figure 6A, right). Our goal was to determine the order of operations most consistent with the joint rod-cone responses. Specifically, we compared a model in which rod and cone signals are combined prior to rectification (Figure 6E right: *common NonLinearity model*), to a model in which rod and cone signals are combined after rectification (e.g. in the ganglion cell; Figure 6E left: *independent NL model*). In the common NL model, the filtered rod and cone signals are scaled, linearly summed, and passed through a common static nonlinearity taken as the average of the independently fit rod and cone nonlinearities (see Methods). In the independent NL model, the filtered signals are transformed separately by their respective nonlinear functions, and then linearly summed. No additional parameters are varied to optimize model predictions in either model. The output of either model is a time-varying signal in units of synaptic current.

To test the two model architectures, uncorrelated rod and cone noise stimuli were presented simultaneously to the retina (Figure 6A, right). Figure 6F compares predicted responses from the common and independent nonlinearity models with the empirically-measured response. We quantified this comparison using the explained variance (Figure 6G). The common nonlinearity model outperformed the independent nonlinearity model in all cases. These findings corroborate the findings from the voltage clamp recordings in Figure 5, as well as previous work (Grimes et al., 2015), indicating that at least some rod and cone signal integration occurs prior to a shared nonlinear circuit element.

We used the model to identify stimuli in which the pre-synaptic integration of rod and cone signals likely shapes ganglion cell inputs the most. To do this, we organized the simultaneous noise stimuli by the value of their rod and cone generator signals (i.e. stimuli passed through the corresponding linear filters) and compared the performance of the two models in this space (Supplementary Figure 3). The independent NL model failed particularly for stimuli that elicited anti-correlated rod- and cone-mediated responses. A similar approach should allow identification of other stimuli that are predicted to lead to weak or strong rod-cone interactions (see Discussion).

### Predictive model correctly predicts response kinetics and rod-cone flicker interference

We next tested whether the model developed in Figure 6 can correctly predict the appropriate delays (Figure 4) and interference between rod and cone flicker that shapes both human perception (Figure 1) and retinal outputs (Figure 2). Revisiting this specific stimulus using the model allowed us to test the consistency of our behavioral, mechanistic, and computational findings. Like direct recordings of excitatory synaptic input to On Parasol RGCs, the output of the common nonlinearity model predicts a high degree of suppression when rod and cone stimuli were presented together at 8 Hz (Figure 7A; measured traces repeated from Figure 5). The model also accurately predicts the phase shifts between rod and cone responses (Figure 7B) across frequency and the absolute amplitude of the rod-cone interactions (Figure 7C) at 8 Hz. Central to the success of the model is the large difference in kinetics of the linear filters for rod and cone stimuli (Figure 7B, inset) and the summation of rod- and cone-mediated responses prior to a shared nonlinearity (Figure 6E, right).

**Figure 7.**
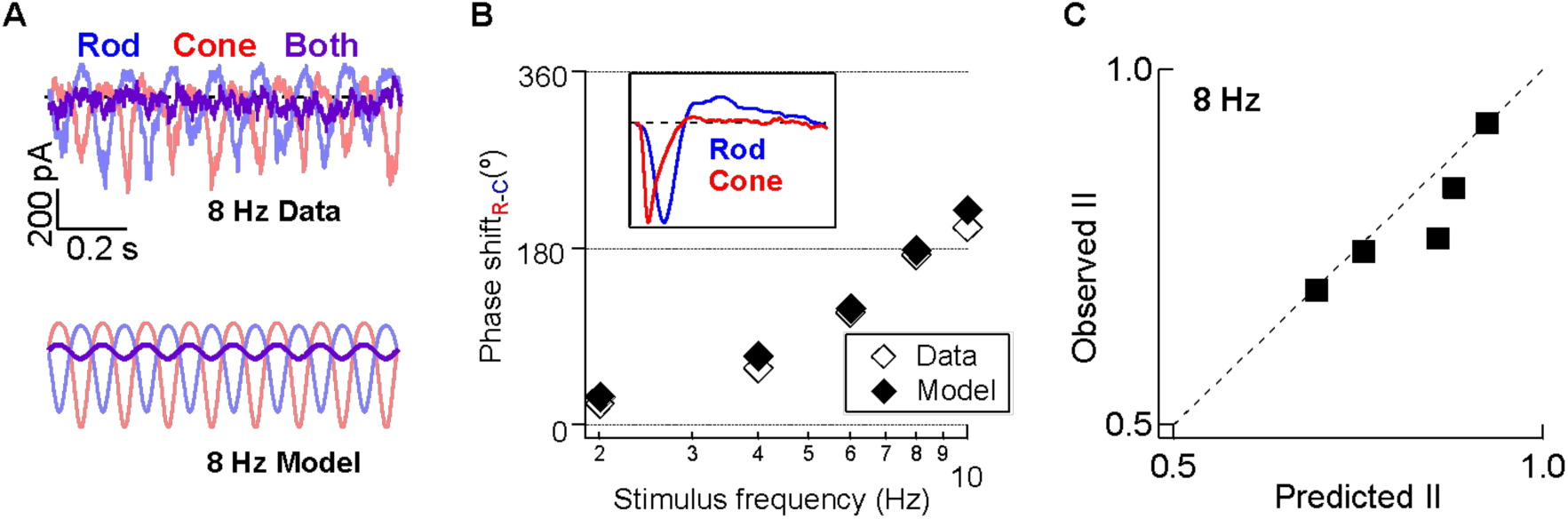
Computational model accurately predicts rod-cone retinal interference in excitatory synaptic input. A) *top* Recording of excitatory synaptic inputs to rod, cone, and combined flicker at 8 Hz. *Bottom* Modelled responses to rod, cone and combined flicker at 8 Hz for the same cell. B) The rod-cone kinetic model captures the frequency-dependence of rod-cone signal delays (i.e. phase shifts. Inset shows linear filters from gaussian noise stimuli, as in Figure 6. C) The rod-cone kinetic model accurately predicts the amplitude of the empirically-measured rod-cone nonlinear interaction index from the same On Parasol RGCs.

## Discussion

Vision at light levels between moonlight and dawn relies on a combination of signals generated by rod and cone photoreceptors. Under these conditions, interactions between rod- and cone-mediated signals shape many aspects of visual perception (reviewed by (Buck, 2004; Buck, 2014; Grimes et al., 2018b)). These interactions are likely to begin within the retina since rod and cone signals converge within the retinal circuitry to modulate signals prior to transmission down the optic nerve (Gouras & Link, 1966; Enroth-Cugell et al., 1977). The circuits conveying rod and cone signals through the retina, including potential sites of interaction, are well known. Here we exploit these properties of mesopic vision to show how common features of parallel processing in neural circuits can explain a perceptual insensitivity to high-contrast flickering lights that modulate both rod and cone signals (MacLeod, 1972).

### Linking the mechanisms controlling parallel neural processing to perception

Interactions between rod and cone signals shape the chromatic, spatial and temporal sensitivity of human perception (reviewed by (Buck, 2004; Buck, 2014; Grimes et al., 2018b)). While the importance of these interactions for how we see has been appreciated for many years, several issues have made it challenging to identify the mechanistic basis of these interactions. First, many rod-cone interactions likely involve both retinal and cortical circuits, and isolating the contributions of each circuit can be difficult (Grimes et al., 2015). Second, it is difficult to generalize between rod-cone interactions in primates and rodents due to differences in visually-guided behavior, in the architecture and cellular composition of retinal circuits, and in the routing of signals through those circuits (Grimes, Baudin, Azevedo, & Rieke, 2018a). Third, the smoothness of visual perception often obscures the complexity of the operation of the underlying circuits - e.g. despite constant involuntary eye movements, we perceive the world steadily.

Here, we investigated the mechanistic substrate for a specific break in the seamlessness of visual perception: a surprising insensitivity to high-contrast flickering lights that activate both rod and cone photoreceptors (MacLeod, 1972). MacLeod suggested that this perceptual insensitivity originated from destructive interference of rod and cone signals in retinal circuits. Our findings confirm this suggestion and identify the retinal circuits and mechanisms within these circuits responsible. We find that destructive interference of rod and cone signals in human perception and in the retinal output share a similar dependence on temporal frequency and phase shift between rod and cone stimuli. These features could be explained by differences in the kinetics of the parallel circuits conveying rod and cone signals through the retina and the convergence of these signals prior to a shared nonlinearity. This provides a clear link between the mechanisms shaping parallel processing and the control of perceptually-relevant circuit outputs.

Several features allowed us to make a tight connection between retinal signaling and perception. First, the perceptual insensitivity to flicker suggests specific kinetic differences between rod and cone signals. Second, the relevant interactions between these kinetically distinct signals likely occur within the retina since rod and cone signals are combined prior to the retinal output. And third, understanding of these kinetic differences makes clear and testable predictions about how specific stimulus manipulations will impact perception. Other perceptual phenomena sharing some of these features - such as independence of adaptation in different photoreceptor types (Chichilnisky & Wandell, 1995; Lee, Dacey, Smith, & Pokorny, 1999) - provide appealing targets for similar attempts to link circuit mechanisms, function and perception.

### Dynamic temporal delays

Interpretation of psychophysical studies of rod-cone interactions often relies on assumptions about how rod signals traverse the retina. A common assumption is that rod signals traverse the retina through different circuits at different light levels. Specifically, the assumption is that the rod bipolar pathway dominates at low mesopic light levels and the (assumed faster) rod-cone pathway, mediated by gap junctions between rods and cones, contributes substantially at high mesopic light levels (Figure 2). Our recent work (Grimes et al., 2018a) suggests that this is not the case, and that instead rod signals traverse the primate retina primarily through the rod bipolar pathway across light levels.

Rod-cone perceptual interactions are also often interpreted with the assumption that, at a given light level, rod signals are delayed relative to cone signals by a fixed amount across stimulus frequencies. Our data is inconsistent with this fixed-delay assumption (Figure 4). Instead, the delay between rod and cone signals depends on temporal frequency; the conserved routing of rod signals suggests that this frequency dependence originates from mechanisms such as inhibition or synaptic depression within the rod bipolar pathway. The apparent differences between the mechanistic operation of retinal circuits under mesopic conditions and the assumptions often used to link these circuits to human perception will require re-evaluating the mechanistic basis of established psychophysical findings.

### Predictive model

Rod-cone interactions are typically studied using highly unnatural stimuli, and understanding rod-cone interactions in natural vision will require exploring a much larger stimulus set. This process can be made efficient by using predictive, empirically-derived models rooted in known mechanisms to explore the role, or competing roles, specific circuit mechanisms play in processing. These predictions can then be tested experimentally, and discrepancies between data and model can shed light on weaknesses in the model architecture and/or reveal previously unknown mechanisms.

With this long-term goal in mind, we sought to develop a predictive model that captured the destructive interference between rod and cone signals. The current instantiation of our model focuses on predicting an On ganglion cell’s excitatory synaptic inputs, particularly the kinetic properties of those inputs. We found that differences in kinetics of rod and cone signals and a shared nonlinearity operating after the signals were combined could account for destructive interference. We hope that extensions of the model to include inhibitory circuits, receptive field subunits and spike generation will allow identification of other stimuli for which rod-cone signal interactions might play an important role in shaping retinal outputs and perception.

## Methods

### Human Psychophysics

The human psychophysics apparatus consisted of one 60 Hz LCD computer monitor (1920 × 1200 Dell, model U2412M) controlled by a Mac mini computer running Psychtoolbox for Matlab (Brainard, 1997; Pelli, 1997). NDF0.6, ‘Bright pink’, and ‘Scarlet’ gel filters (Rosco Ecolour, Stamford, CT) were mounted to the front of the monitor to control luminance and suppress wavelengths between 500–600 nm, thus improving the photoreceptor selectivity of the red and blue phosphors. Human observers fixated a small cross while red (peak power at 640 nm; L-cone-preferring) and blue (peak power at 444 nm; rod-preferring) 2° spots were presented at ∼10° eccentricity to the observer’s retina.

In the rod equivalent matching task, additional NDF2 and NDF0.5 filters were added. Dark adaptation time was 30 minutes. There were three subtasks: (1) finding the rod threshold, (2) finding the L-cone threshold, (3) determine the rod activity associated with the red LED (e.g. the rod equivalent match). In the first subtask, observers adjusted the intensity (measured in R*/L-cone/second) of a red spot in their periphery until it was barely detectable. In the second subtask, observers adjusted the intensity of a red patch until hue was barely detectable. In the third subtask, the observer was shown a fixed intensity red patch for 800ms, then an adjustable blue patch for 600ms, then the same initial fixed intensity red patch again for 600ms. For this subtask the observer adjusted the intensity of the blue spot until it matched the apparent intensity red signal. The intensity of the fixed red spot was kept at 75% under the cone (hue) threshold and 25% above the rod (patch detection) threshold. Based on this unique measurement of rod activity elicited by the red spot for each observer we were able to improve the selectivity of the red flash for L-cone activation by using temporally-matched proportional decrements in the blue mean (i.e. silent substitution).

In the flicker detection threshold task, the NDF2 and NDF0.5 filters were removed. Dark adaptation time for these experiments was 20 minutes. Observers adjusted the contrast of a flickering spot between 5-95% (decreasing the contrast if the flicker was detectable, or increasing the intensity if the flicker was not detected) until they had reversed the direction of their adjustment (referred to as a ‘crossing’) 8 times. From these crossing values we calculated the threshold from the weighted averages between crossings. The combined mean was kept constant at 2 R*/rod/s,. After achieving a threshold the task automatically advanced to the next trial, 4-6 trials were conducted for each condition and weighted thresholds were averaged across trials. An adaptive algorithm was included to speed the time required to find an observer’s threshold; the size of the contrast adjustments were decreased after every crossing. To minimize onset effects, the patch was initially presented statically at the mean luminance for 0.5 seconds, followed by 2.5 s of sinusoidal flicker at an adjustable contrast.

There were two types of flicker experiments, the first explored rod-cone interactions across temporal frequencies and the second tested the effect of temporal delays (i.e. phase shifts) at two frequencies (4 and 8 Hz). In the first experiment, we tested four frequencies (4, 6.5, 8, 9.5 Hz), with three conditions (L-cone targeted flicker only, rod-targeted flicker only, then rod-cone combined flicker) at each frequency. Values obtained from the independent presentation of rod- or cone-preferring stimuli were used to fix the contrast ratio for rod and cone stimuli on trials with combined presentation. In the second experiment, the observer’s task was identical. Again, the ratio between the rod and cone thresholds was determined from independent stimuli presentation at a single frequency (8 Hz). Cone-preferring stimuli were then temporally offset (from rod-preferring stimuli) during combined presentations by phase shifts of 0º, 45º, 90º, 135º, and 180º.

### Tissue preparation and storage

Non-human primate retina was obtained through the Tissue Distribution Program of the Regional Primate Research Center at the University of Washington. Experiments were conducted on whole mount preparations of isolated primate retina as previously described (Dunn, Lankheet, & Rieke, 2007; Trong & Rieke, 2008). In brief, pieces of retina attached to the pigment epithelium were stored in ∼32–34°C oxygenated (95% O_2_/5% CO_2_) Ames medium (Sigma, St Louis, MO) and dark-adapted for >1 hr. Pieces of retina were then isolated from the pigment epithelium under infrared illumination and flattened onto polyL-lysine slides. Once under the microscope, tissue was perfused with oxygenated Ames medium at a rate of ∼8 ml/min.

### Electrophysiology

Extracellular recordings from ON and OFF Parasol retinal ganglion cells were conducted using ∼3 MΩ electrodes containing Ames medium. Voltage-clamp whole-cell recordings were conducted with electrodes (3–4 MΩ) containing (in mM): 105 Cs methanesulfonate, 10 TEA-Cl, 20 HEPES, 10 EGTA, 2 QX-314, 5 Mg-ATP, 0.5 Tris-GTP and 0.1 Alexa (488, 555 or 750) hydrazide (∼280 mOsm; pH ∼7.3 with CsOH). Current-clamp whole-cell recordings from horizontal cells were conducted with (5–6 MΩ) electrodes containing (in mM): 123 K-aspartate, 10 KCl, 10 HEPES, 1 MgCl_2_, 1 CaCl_2_, 2 EGTA, 4 Mg-ATP, 0.5 Tris-GTP and 0.1 Alexa (488, 555 or 750) hydrazide (∼280 mOsm; pH ∼7.2 with KOH). In initial experiments, cell types were confirmed by fluorescence imaging following recording. To isolate excitatory or inhibitory synaptic input, cells were held at the estimated reversal potential for inhibitory or excitatory input of ∼−60 mV and ∼+10 mV. These voltages were adjusted for each cell to maximize isolation. Absolute voltage values have not been corrected for liquid junction potentials (K^+^-based = −10.8 mV; Cs^+^-based = −8.5 mV).

Visual stimuli (diameter: 500–560 μm) were delivered to the preparation through a customized condenser from blue (peak power at 460 nm) or red (peak power at 640 nm) LEDs. Light intensities (photons/μm^2^/s) were converted to photoisomerization rates (R*/photoreceptor/s) using the estimated collecting area of rods and cones (1 and 0.37 μm^2^, respectively), the stimulus (i.e., LED or monitor) emission spectra and the photoreceptor absorption spectra (Baylor, Nunn, & Schnapf, 1984; Baylor, Nunn, & Schnapf, 1987). The blue and red LEDs produced a mean of ∼20 R*/rod/s and ∼200 R*/L-cone/s. Rod- and cone-preferring flashes were 10 ms in duration.

### Modeling

Components of the Linear-nonlinear Cascade model (Figure 6) were derived from voltage-clamp recordings of full-field white noise (0-40 Hz bandwidth) using the long and short wavelength LEDs as previously described (Chichilnisky, 2001; Grimes et al., 2015). Model components were verified by taking the model’s explained variance from data with the same noise stimuli presented. Model predictions to the sine wave stimuli were a result of a three-stage process: 1) rod and cone stimuli are convolved with the respective linear filters 2) filtered signals are summed and 3) combined signal is passed through the average nonlinearity. The continuous output signals are in units of picoAmps.

The vector summation interference model (Figure 4) reflects a linear summation of rod and cone signal amplitudes scaled by the degree to which the signals are in phase (i.e. cosine function). Signal interactions in this model depend on stimulus frequency and the time delays and amplitudes of the responses to individually-delivered stimuli.

## Acknowledgements

We thank Jeff Diamond, Belle Liu, Arthur Hung and Mike Manookin for helpful comments on an earlier version of the manuscript, and Shellee Cunnington and Chris English for excellent technical support. Funding was provided by the NIH (EY028111 and 5R90DA033461).

---

**Supplementary Figure 1 (related to Figure 1).**
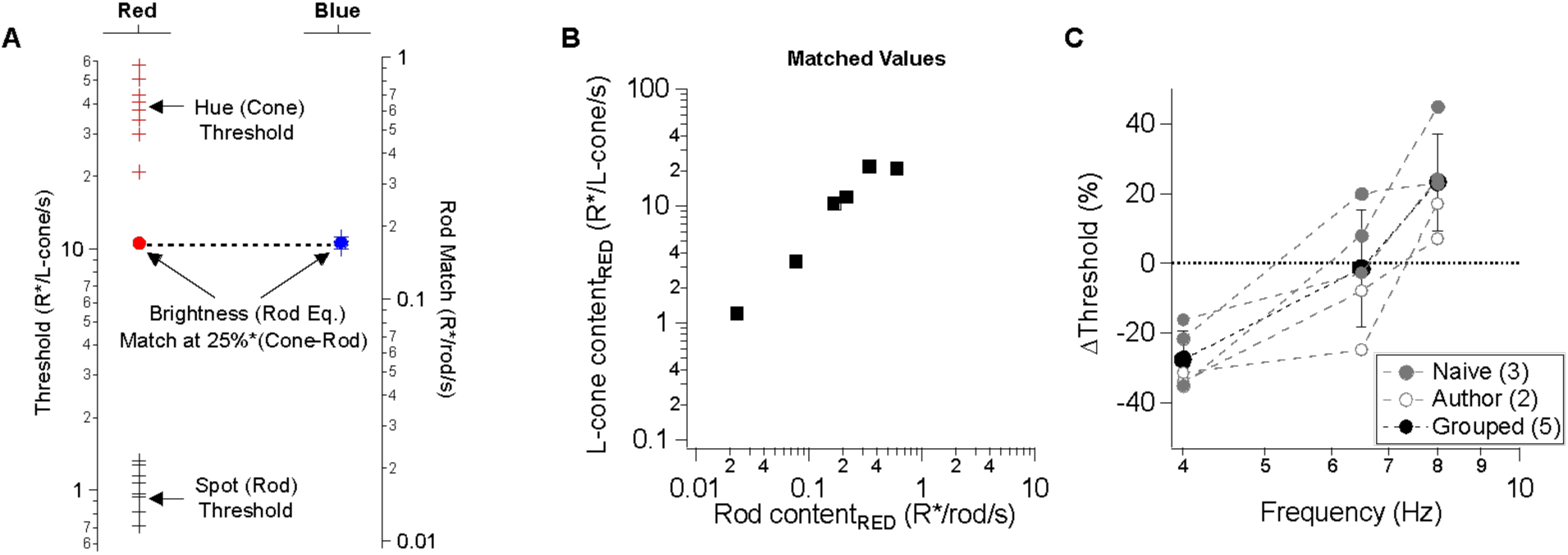
Additional details pertaining to the psychophysical experiments. A) To improve the selectivity of the L-cone-preferring stimulus, we obtained rod-equivalent matches for each observer under scotopic conditions and utilized the results to create silent-substitution under mesopic conditions (see Methods). An observer first finds the detection threshold for a static long wavelength spot. Next the observer finds a hue threshold for the same spot. Lastly, the long wavelength spot is fixed at 25% of the difference between the average hue and detection thresholds and another short wavelength spot is adjusted until a brightness match is achieved. B) Rod-equivalent matches for all observers. Plot depicts the empirically measured rod content of the long wavelength spot. C) Changes in flicker thresholds for combined stimulus presentations for each observer (i.e. breakdown of Figure 1C). A) Spike responses to cone (*top*, red), rod (*middle*, blue), or combined rod-cone (*bottom*, purple) flicker (2-10 Hz) in an On Parasol retinal ganglion cell. A strong suppressive interaction was observed when rod and cone stimuli were flickered together at 8 Hz. B-C) Mean spike responses (sinusoidal fit, see Methods) as a function of stimulus frequency for 6 On Parasol RGC recordings (B) and 5 Off Parasol RGC recordings (C). D-E) Relative timing differences (i.e phase shifts) between isolated rod and cone responses for On (D) and Off (E) Parasol RGCs. Markers and error bars in B-E represents mean±SEM.

**Supplementary Figure 2 (related to Figure 3):**
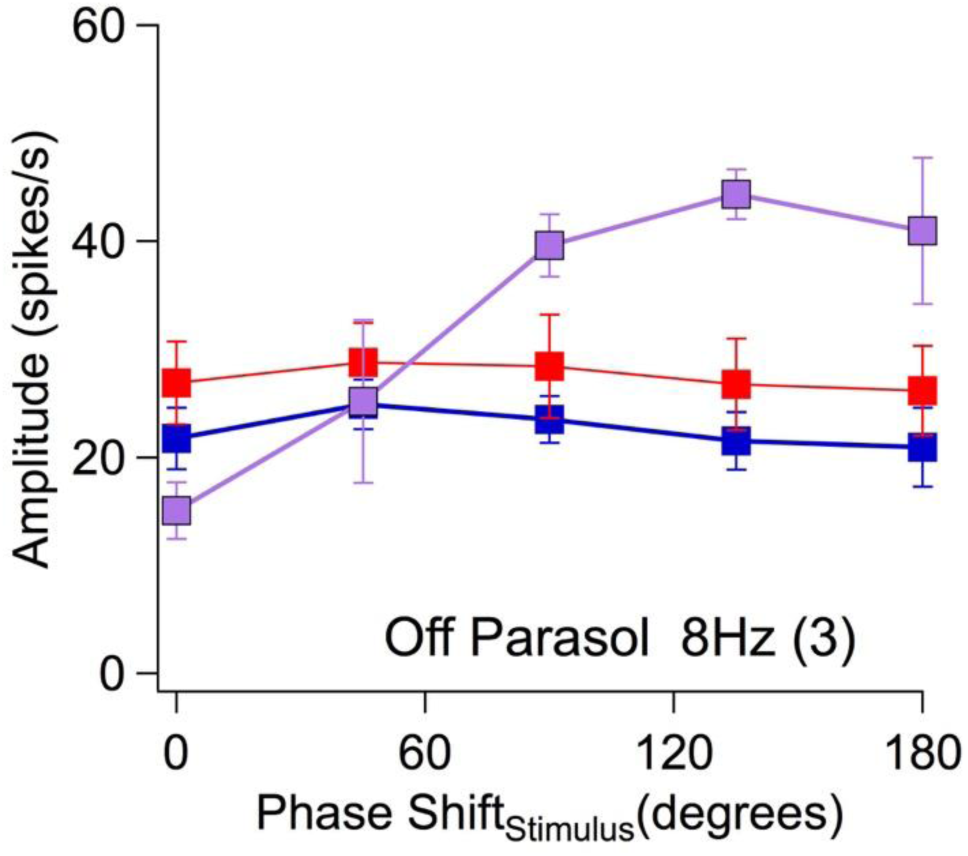
Rod-cone signals constructively and destructively interfere within Off Parasol RGC circuits. At 8 Hz, rod and cone signals can be destructive or constructive depending on the relative delay between the rod and cone stimuli.

**Supplementary Figure 3 (related to Figure 6):**
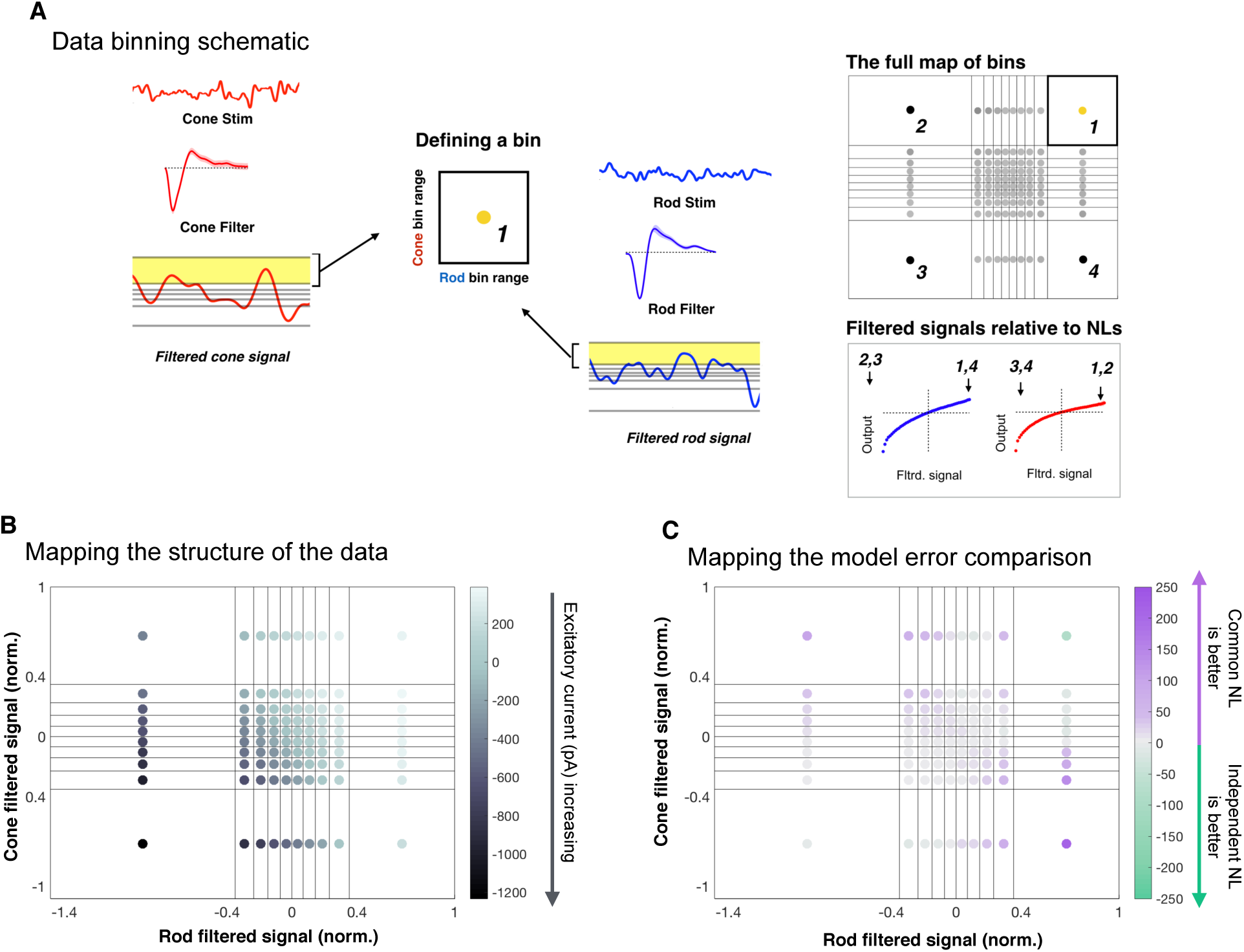
Dependence of model performance on specific rod-cone contributions to the signal. A) Regardless of the model architecture (e.g. common or independent nonlinearities), the initial linear filtering stage is separate for the rod and cone branches of the model. Hence, we sort the data based on bins based on the rod and cone filtered signals. The bins are equally populated with data points across five cells. A) A diagram representing the horizontal and vertical ranges of each bin (note: diagram does not depict real bin divisions). B) In plotting the mean current value across single bins, we see a clear underlying structure (i.e. excitatory current input increases as both filtered signals become both larger and more negative). This provides assurance that the binning method cleanly captures the data. C) The error between a model and the data in a single bin is taken as the absolute difference between the mean data and the mean model output. We compare the errors of the two models by simply subtracting the common nonlinearity error from the independent nonlinearity error (as the independent model had the larger average error). The common nonlinearity model outperforms the independent model in nearly every case where there is a difference in performance difference between the two models; in particular, this improvement was largest along the diagonal where the rod and cone filtered stimuli are the most anti-correlated.

